# A Single-Cell RNA-Seq Analysis Unravels the Heterogeniety of Primary Cultured Human Corneal Endothelial Cells

**DOI:** 10.1101/2023.02.09.527872

**Authors:** Pere Català, Nathalie Groen, Vanessa L.S. LaPointe, Mor M. Dickman

**Affiliations:** Department of Cell Biology–Inspired Tissue Engineering, MERLN Institute for Technology-Inspired Regenerative Medicine, Maastricht, the Netherlands; University Eye Clinic Maastricht, Maastricht University Medical Center+, Maastricht, the Netherlands; Single Cell Discoveries, Utrecht, the Netherlands

## Abstract

The primary culture of donor-derived human corneal endothelial cells (CECs) is a promising cell therapy. It confers the potential to treat multiple patients from a single donor, alleviating the global donor shortage. Nevertheless, this approach has limitations preventing its adoption, particularly culture protocols allow limited expansion of CECs and there is a lack of clear parameters to identify therapy-grade CECs. To address this limitation, a better understanding of the molecular changes arising from the primary culture of CECs is required. Using single-cell RNA sequencing on primary cultured CECs, we identify their variable transcriptomic fingerprint at the single cell level, provide a pseudo temporal reconstruction of the changes arising from primary culture, and suggest markers to assess the quality of primary CEC cultures. This research depicts a deep transcriptomic understanding of the cellular heterogeneity arising from the primary expansion of CECs and sets the basis for further improvement of culture protocols and therapies.

## INTRODUCTION

The cornea is the transparent window transmitting light into the eye. The inner part of this avascular tissue is covered by a monolayer of hexagonal corneal endothelial cells (CECs) (DelMonte and Kim, 2011) that maintain corneal transparency and hydration by their pump and barrier function (Bonanno, 2012). Human CECs are arrested in a non-proliferative state and lack regenerative capacity. Consequently, damage to CECs due to surgery, inherited diseases, or acquired conditions results in irreversible corneal oedema, impairing vision. (Price et al., 2021).

Corneal transplantation is the current therapy for treating corneal endothelium dysfunction. Still, only one donor cornea is available for every 70 patients in need, leaving 12.7 million people awaiting treatment worldwide (Gain et al., 2016). A first landmark clinical trial showed that primary cultivated CECs can restore corneal transparency, breaking the one-donor–one- recipient paradigm. (Kinoshita et al., 2018; Peh et al., 2015a). Encouraged by the long-term success of this therapy, (Numa et al., 2021) clinical trials are ongoing in Japan (UMIN000034334 and UMIN000012534), Mexico (NCT04191629) and Singapore (NCT04319848) to assess the therapeutic potential of cultured CECs.

Nonetheless, the transplantation of cultured CECs has limitations preventing its wider adoption. Primary CEC cultures are only successful when derived from donors younger than 45 years of age, limiting the pool of donor corneas suitable for this technique. Furthermore, cultures become heterogeneous over time, and significant alterations diminishing the cell phenotype and functionality are observed after the second passage (Frausto et al., 2020). Notably, there is a lack of clear parameters to identify therapy-grade cells (Català et al., 2022). Recently, cell morphology (Yamamoto et al., 2019) and a set of markers: CD44, CD105, CD24, and CD133, also referred to as the E-ratio, have been used as exclusion criteria for therapy-grade CECs (Ueno et al., 2022). If we were able to identify additional or other cell-specific markers we could selectively assess and enrich for therapy-grade CECs.

To deconstruct the heterogeneity and gain knowledge on the alterations arising from the primary culture of CECs, we used single-cell RNA sequencing (scRNA-Seq) to profile 42,220 primary human CECs from six corneas of three donors at five time points over three passages in culture. Our analysis revealed that the culture diversified over time into heterogeneous subpopulations including cells less desirable for therapy that were entering a senescent or fibrotic state. We identified markers that can be used in combination to assess for therapy-grade cells and enrich for desired cell populations. Pseudo time analysis further uncovered the different trajectories arising during culture. Together, our study sheds light on the various routes followed by CECs in culture, identifies novel markers to increase culture efficiency, and presents a roadmap to improve culture protocols.

## RESULTS

### scRNAseq reveals different subpopulations of CECs arising from primary culture

Six paired human corneas, from two male and one female donor (Table 1), were used for isolation, primary culture, and scRNAseq of CECs (Figure 1A). Cells from five different time points in culture of were loaded for sequencing as specified in Table 2. Namely, cells at days 2 and 5 of culture after their isolation in M4 proliferation media at passage 0, and cells at passages 0, 1, and 2, at confluency after 7 days of culture in M5 stabilization media. The cultures showed characteristic CEC morphology over time (Figure 1B, Figure S1) and expressed the desired CEC proteins such as CD166 and zonula occludens-1 (ZO-1) (Figure 1c).

**Figure 1.**
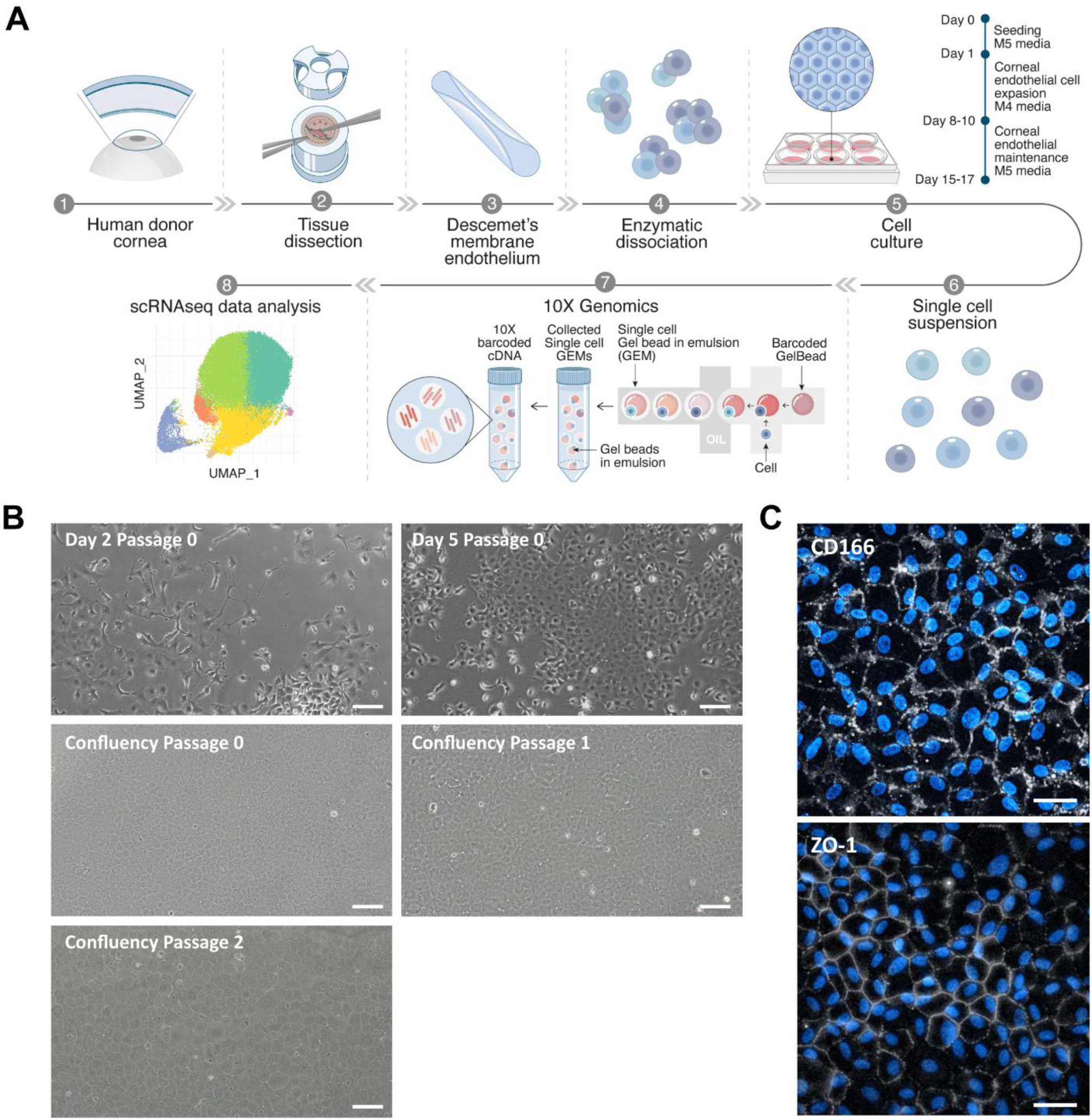
Human CECs were successfully isolated and cultured for scRNAseq. (A) Schematic representation of the experimental overview. (B) Phase contrast images of CEC over the different time points selected for scRNAseq confirm the desired hexagonal morphology of the cells. Scale bars represent 100 μm. (C) Immunofluorescence assessment of CEC markers CD166 and ZO-1 confirm the phenotype of the primary cultured cells at passage 2. Scale bars represent 50 μm.

**Table 1.** Donor cornea information. COD= cause of death, GI= Gastrointestinal, MVA= motor vehicle accident, ECD= endothelial cell density, OS= oculus sinister, OD= oculus dexter.

| Donor | Sex | Age (years) | Preservation time<br>(days) | COD | ECD OS/OD (cells/mm<sup>2</sup><br>) |
| --- | --- | --- | --- | --- | --- |
| Donor 1 | Male | 34 | 12 | Trauma | 3195/2917 |
| Donor 2 | Male | 29 | 11 | GI bleed | 2832/2878 |
| Donor 3 | Female | 27 | 14 | MVA | 2861/3159 |

**Table 2.**
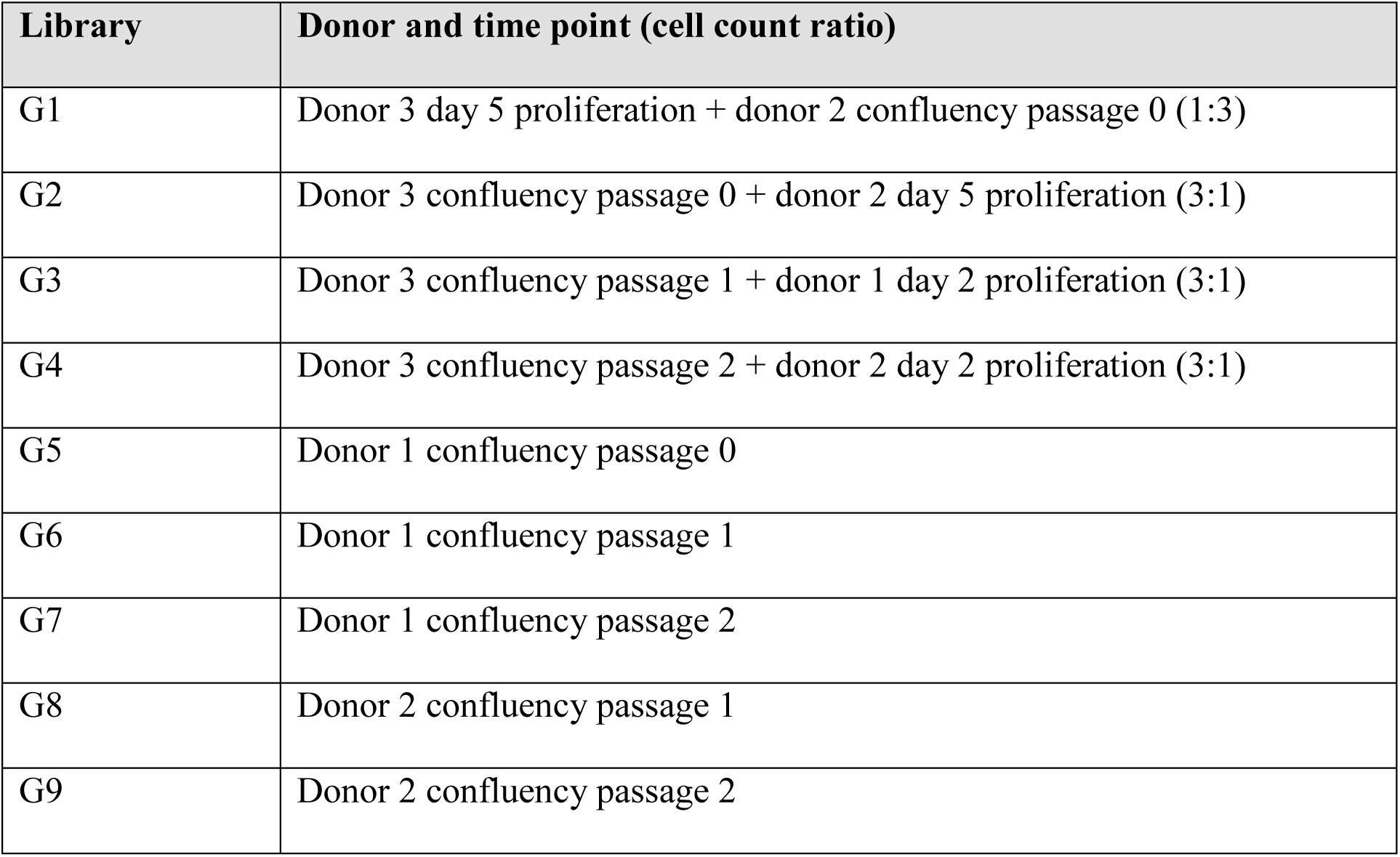
10x genomics sample loading and library information.

| Library | Donor and time point (cell count ratio) |
| --- | --- |
| G1 | Donor 3 day 5 proliferation + donor 2 confluency passage 0 (1:3) |
| G2 | Donor 3 confluency passage 0 + donor 2 day 5 proliferation (3:1) |
| G3 | Donor 3 confluency passage 1 + donor 1 day 2 proliferation (3:1) |
| G4 | Donor 3 confluency passage 2 + donor 2 day 2 proliferation (3:1) |
| G5 | Donor 1 confluency passage 0 |
| G6 | Donor 1 confluency passage 1 |
| G7 | Donor 1 confluency passage 2 |
| G8 | Donor 2 confluency passage 1 |
| G9 | Donor 2 confluency passage 2 |

After filtering for cells with a minimum of 1,000 transcripts, the transcriptome profiles of 42,220 cells were embedded in a uniform manifold approximation and projection (UMAP). Cells from all donors were distributed homogeneously across the UMAP (Figure S2). Unbiased low-resolution clustering revealed six major cell clusters (Figure 2A), all of which expressed CEC markers *ALCAM* (CD166), *PRDX6*, *SLC4A11*, *PITX2*, and *ATP1A1*, (Van den Bogerd et al., 2019; Català et al., 2020, 2022) confirming their endothelial identity (Figure 2B). The absence of keratocyte markers *CD34*, *KERA*, and *ALDH1A1*, (Fernández-Pérez and Ahearne, 2019; Foster et al., 2015) and epithelial markers *KRT12*, *KRT14*, and *PAX6* (Hayashi et al., 2016; Ouyang et al., 2014) confirmed the absence of contaminating cell types from corneal stroma and epithelium (Figure S3). Differential gene expression profiling was used for cluster identification.

**Figure 2.**
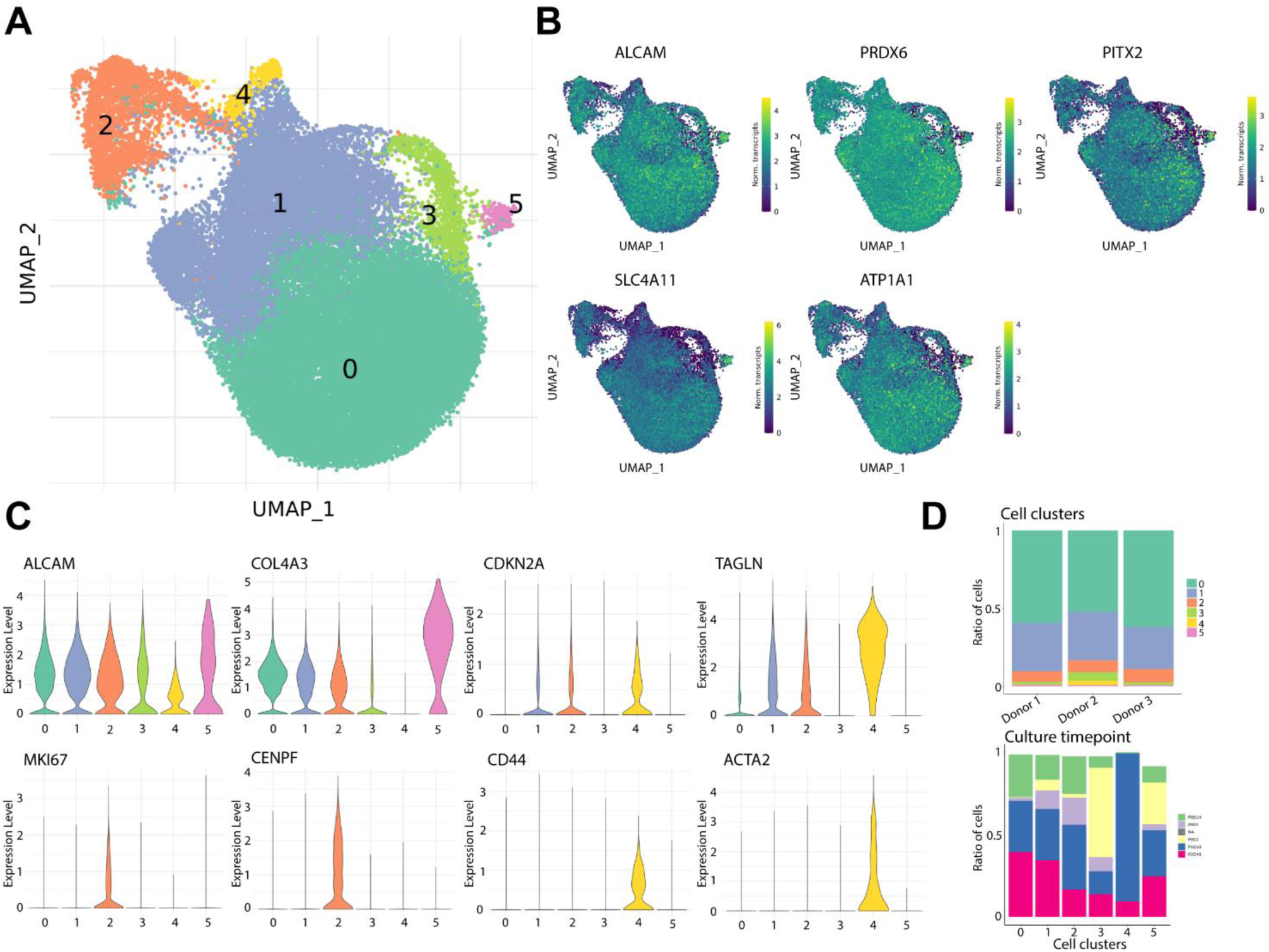
scRNAseq analysis reveals distinct clusters of primary cultured CECs. (A) UMAP of the 42,220 sequenced cells reveals six cell clusters. (B) Gene expression UMAP of typical CEC markers namely *ALCAM* (CD166), *PRDX6*, *PITX2*, *SLC4A11*, and *ATP1A1* confirms the endothelial identity of the sequenced cells. (C) Violin plots of show gene expression for markers of endothelium (*ALCAM*, *COL4A3*), senescence (*CDKN2A*, *TAGLN*), proliferation (*MKI67*, *CENPF*), and fibrosis (*CD44*, *ACTA2*). (D) Bar charts show the distribution of cells of each donor per cluster (top) and the time point composition of each cluster (bottom).

Clusters 0 and 5 presented increased differential gene expression of typical CEC markers, *SLC4A11*, *COL4A3* (Van den Bogerd et al., 2019), *CDH2* (He et al., 2016), and *ALCAM*, compared to the other sequenced cells, suggesting these clusters were composed of therapy-grade CECs (Figure 2C and S4). Cluster 1 was identified as CECs transitioning towards a senescent phenotype due to the high differential expression of senescence markers such as *MT2A* (Malavolta et al., 2017), *CDKN2A* (p16) (Enomoto et al., 2006), and *TAGLN* (Benadjaoud et al., 2022) (Figure 2C and S4). Cluster 2 was composed of highly proliferative CECs expressing *MKI67* (Sun and Kaufman, 2018), *CENPF* (Varis et al., 2006), and *PTTG1* (Ersvær et al., 2020) (Figure 2C). Cluster 4 was composed of fibrotic CECs with increased differential expression of *ACTA2* (α-smooth muscle actin (SMA)) (Rao et al., 2014), *CD44* (Hamuro et al., 2016), and *COL6A1* (Oouchi et al., 2021) (Figure 2C and S4). Finally, our analysis revealed that cluster 3 was mainly composed of cells in passage 0 (at both day 2 and day 5) (Figure 2D) that differentially expressed ribosomal-associated genes (Figure S5). This finding suggested that cells at an early culture stage clustered together due to the necessary adaptation to *in vitro* culture conditions and the use of proliferation media, which led us to further explore the cells that had been cultured to confluency in passages 0, 1 and 2.

### scRNAseq reveals seven distinct subpopulations of primary cultured CECs at therapeutically relevant time points

To identify the meaningful differences at therapeutically relevant time points and remove the clustering bias introduced by the adaptation to primary culture conditions after cell isolation, we separately analyzed the CECs at confluency in passages 0, 1, and 2. These time points are the most therapeutically relevant, as CECs are most suitable for therapy after 7 days in M5 stabilization media up to passage 2(Frausto et al., 2020; Peh et al., 2019).

After removing sub-confluent cells from day 2 and day 5 (in passage 0) of culture, the transcriptome profiles of 37,158 CECs at confluency in passages 0, 1, and 2 were embedded in a UMAP. Unbiased low-resolution clustering revealed seven cell clusters (Figure 3A) with distinct transcriptomic signatures. Cells from all donors were distributed homogeneously across the UMAP (Figure S6). Differential gene expression analysis was used for identification of each cell cluster (C0–C6).

**Figure 3.**
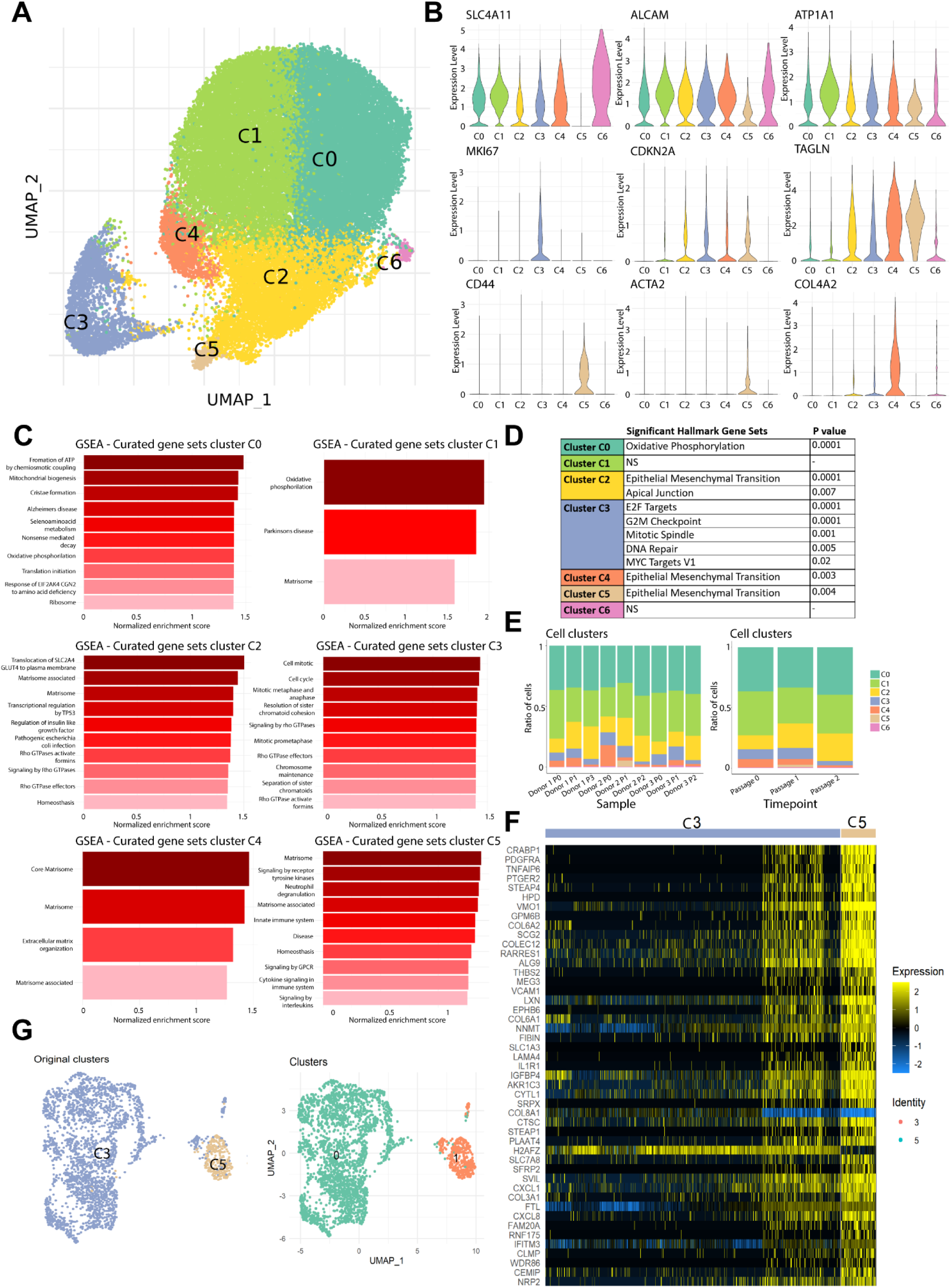
scRNAseq analysis reveals seven distinct CEC clusters at therapeutically relevant time points. (A) UMAP of the 37,158 cells at confluency time points reveals seven cell clusters. (B) Violin plots show gene expression for markers of endothelium (*SLC4A11*, *ALCAM*, *ATP1A1*), proliferation (*MKI67*), senescence (*CDKN2A*, *TAGLN*), fibrosis (*CD44*, *ACTA2*), and extracellular matrix production (*COL4A2*). (C) Gene set enrichment analysis (GSEA) reveal differentially expressed gene sets across cell clusters (p <0.05). (D) Significant differentially expressed hallmark gene sets across clusters (p <0.05) show endothelial to mesenchymal transition in lower quality CEC clusters C2, C4 and C5; and proliferation hallmarks in cluster C3. (E) Bar charts show composition of cell clusters across the different sequenced samples (left), and across the different time points (right). (F) Heatmap of the top 50 differentially expressed genes across clusters C3 and C5 (p <0.01) shows a subpopulation of CEC within cluster C3 that highly resembles the cells comprising cluster C5. (G) UMAP of the reclustering analysis of cell clusters C3 and C5 with original clusters (left) and newly detected clusters (right).

Increased differential expression of typical CEC markers, such as *COL4A6* (Van den Bogerd et al., 2019), *SLC4A11*, *ATP1A1*, *COL4A3*, and *CDH2* suggested that clusters C0 and C1 were composed of therapy-grade CECs (Figure 3B and S7). Further analysis by gene set enrichment analysis (GSEA) revealed that highly metabolically active cells comprised these clusters (Figure 3C), with a distinct hallmark for oxidative phosphorylation in cluster C0 (Figure 3D). These markers are also found in native functioning human CECs, suggesting these cells were therapy-grade CECs. Cluster C3 was composed of proliferating CECs with differential high expression of *MKI67* (KI-67), *CENPF*, and *PTTG1* (Figure 3B and S7). GSEA further confirmed enriched gene sets and significant hallmarks related to cell proliferation (Figure 3C, D).

Clusters C2 and C4 were composed of CECs with increased differential expression of senescence related genes. Namely, *CDKN2A* (p16), *TAGLN*, and *MT2A* (Figure 3B and S7) in cluster C2 and *CDKN1A* (p21) (Matthaei et al., 2012), *CDKN2A* (p16), and *TAGLN* in cluster C4 (Figure 3B and S7), suggesting these cells were transitioning towards an undesirable senescent and fibrotic phenotype. Cluster C4 had a high extracellular matrix production suggested by the differential high expression of *COL4A1*, *COL4A2*, *COL5A1*, and *FBLN5* in cells that maintained the expression of endothelial markers such as *SLC4A11*, and *COL4A3* (Figure 3B and S7). In contrast, cells in cluster C2 presented a low expression of CEC markers such as *ALCAM*, *SLC4A11*, *CDH2*, and *ATP1A1* (Figure 3B and S7). GSEA revealed that both clusters C2 and C4 had a significant hallmark for epithelial to mesenchymal transition (Figure 3D), confirming these cells were transitioning towards senescence and fibrosis. GSEA revealed that cluster C2 was enriched for genes related to alterations in the matrisome production, upregulation of p53 pathway, and upregulation on Rho GTPase pathway, which are known to regulate cellular senescence (Orgaz et al., 2014; Rufini et al., 2013) (Figure 3C), and cluster C4 expression was enriched for genes related to matrisome, extracellular matrix organization and matrisome associated genes suggesting a remodeling of the extracellular matrix (Figure 3C). Cluster C5 was composed of fibrotic cells differentially expressing the fibrosis-associated markers *COL6A1*, *COL6A3*, *CD44*, and *ACTA2* (Figure 3B). The cells in cluster C5 had diminished expression of *ALCAM* (CD166) and lacked *SLC4A11* expression, two CEC markers. GSEA further suggested enriched expression of genes related to the matrisome and matrisome associated processes and increased signaling by G-coupled protein receptors and receptor tyrosine kinase (Figure 3C), with a significant hallmark for epithelial to mesenchymal transition for cells in cluster C5 (Figure 3D). Finally, Cluster C6 was composed of CECs with differential high expression of typical endothelial markers such as *SLC4A11*, *COL4A3*, and *CDH2* (Figure 3B) suggesting they were therapy-grade CECs. These cells differentially expressed higher *GOLGA8A* and *GOLGA8B*, suggesting a possible increase in secretory pathways. GSEA did not reveal significant upregulation of gene sets nor significant hallmarks compared to other clusters.

### Longer culture times decrease proliferation and increase transitioning to senescence

Our scRNAseq data analysis revealed that at longer culture time points, namely confluency in passage 2, there was an increase in the number of cells transitioning to a senescent/fibrotic phenotype (cluster C2 and C4) and a decrease in the number of proliferative cells (cluster C3) compared to earlier time points (Figure 3E). Interestingly, the number of cells in cluster C4 decreased over culture time (Figure 3E) while cells in cluster C2 increased over time. This could be because the cells in C4 are early senescent cells and transition to later senescent cells in cluster C2. Regarding fibrotic CECs in cluster C5, these cells were detected as early as passage 1, but were also present in passage 2 at lower prevalence compared to passage 1. Finally, the prevalence ratio of therapy grade CECs (cluster C0 and C1) was maintained over time points (Figure 3E) showing the presence of therapy-grade CECs over all culture passages.

### scRNAseq subclustering analysis revealed two distinct transcriptomic profiles of proliferating cells

The CEC marker *ALCAM*, and the fibrotic markers *CD44* and *ACTA2* were heterogeneously expressed across different cells comprising comprising proliferative cluster C3 (Figure S8), suggesting it contained a subcluster of highly proliferative fibrotic CECs. Correlation and differential expression analysis revealed similarities between a subcluster of C3 with the fibrotic cells present in cluster C5 (Figure 3F). This was confirmed by a reclustering analysis of only clusters C3 and C5, which showed that a small subpopulation of cells originating from cluster C3 clustered together with the cells originating from cluster C5 (Figure 3G). This finding confirms the presence of a subpopulation of highly proliferative fibrotic CECs within cluster C3.

### Primary cultured CECs resemble native human CECs

To assess how comparable primary cultured CECs are to native human CECs, we integrated the transcriptome of cells cultured to confluency in passages 0, 1, and 2 with a previously published cornea scRNAseq atlas (Català et al., 2021). To do so, the cluster information from both the CECs in the cornea cell atlas and this study were overlaid in 2D space (Figure 4A). The clustering analysis revealed that the CEC clusters originating from the native human corneal endothelium (Atlas En0 and Atlas En1) clustered adjacent to primary CEC clusters C6, C1 and C0, suggesting these cell clusters share comparable transcriptomic profiles (Figure 4A).

**Figure 4.**
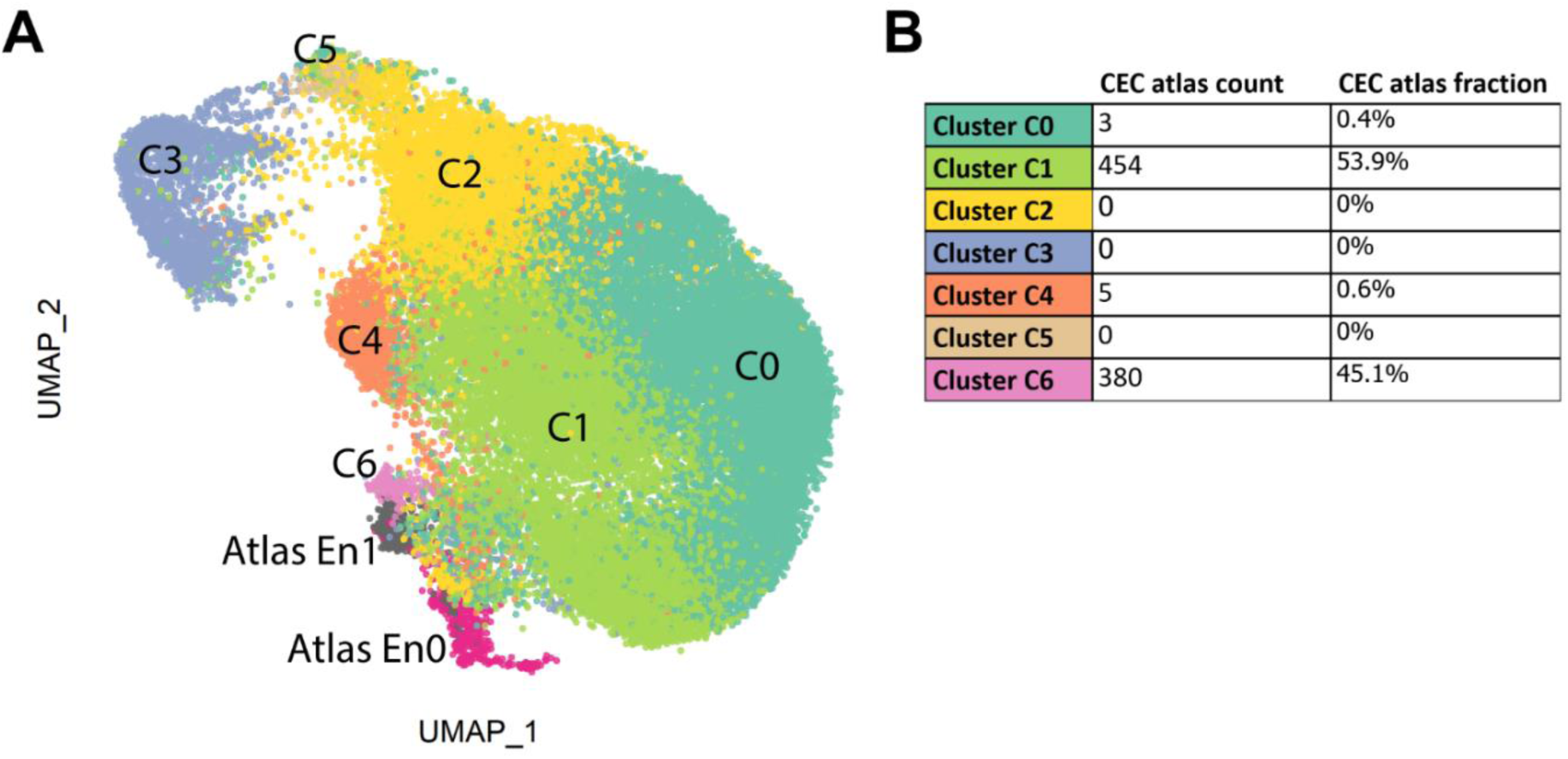
Therapy grade primary cultured CECs resemble human native CECs. (A) Integrated data UMAP of primary cultured CECs at confluency and native human CECs from the previously published cornea cell atlas (Catala et al. 2021). The integration analysis shows that native CECs (Atlas 0 and Atlas 1) cluster adjacent to high quality primary cultured CEC clusters (clusters C0, C1, and C6) suggesting clusters C0, C1 and C6 can be considered therapy-grade primary cultured CECs. (B) The cell type of the atlas CECs was predicted using the CEC confluency dataset using the cell label transfer functionality from Seurat. 99.4% of the atlas cells were attributed to clusters C0, C1 and C6, further suggesting the therapeutic standard of these clusters.

To further understand the similarity of native human CEC to primary cultured cells, we performed a cluster prediction analysis of the atlas CECs using the clustering analysis of cells at confluency. The prediction analysis revealed that 99.4% of the atlas CECs were associated to clusters C1 (53.9%), C6 (45.1%), and C0 (0.4%) (Figure 4B), suggesting these clusters are similar to native human CECs and could be used for therapeutic purposes. Furthermore, our prediction analysis found that no native CECs were associated with the senescent and fibrotic clusters C2 and C5, or the proliferative cluster C3 (Figure 4B).

### Pseudo time reconstruction and evaluation reveal the dynamics of CEC profiles arising from primary expansion

To assess how the cells transition between clusters over time, we performed a pseudo time reconstruction of the cells at confluency in passages 0, 1, and 2. Our first analysis revealed that the CECs originating from clusters C0, C1, and C6, transitioned into the senescent cells in cluster C2, and then became the fibrotic cells in cluster C5 (Figure 5A). Interestingly, the last cluster the pseudo time trajectory identified were the proliferative cells in cluster C3. We hypothesize this is due to the presence of a side population of fibrotic proliferative cells within cluster C3 (Figure 3F, G), which might interfere with the pseudo time trajectory analysis. To reduce such bias and identify gene trends over the pseudo time trajectory, we performed a pseudo time analysis of the confluent cells excluding cluster C3 (Figure 5B). In line with the first analysis, the pseudo time trajectory revealed that the CECs from clusters C0, C1, and C6 transitioned into the transitioning senescent cells in cluster C2, and then the fibrotic cells in cluster C5 (Figure 5B) showing that primary cultured CECs transition towards senescence and fibrosis over culture time.

**Figure 5.**
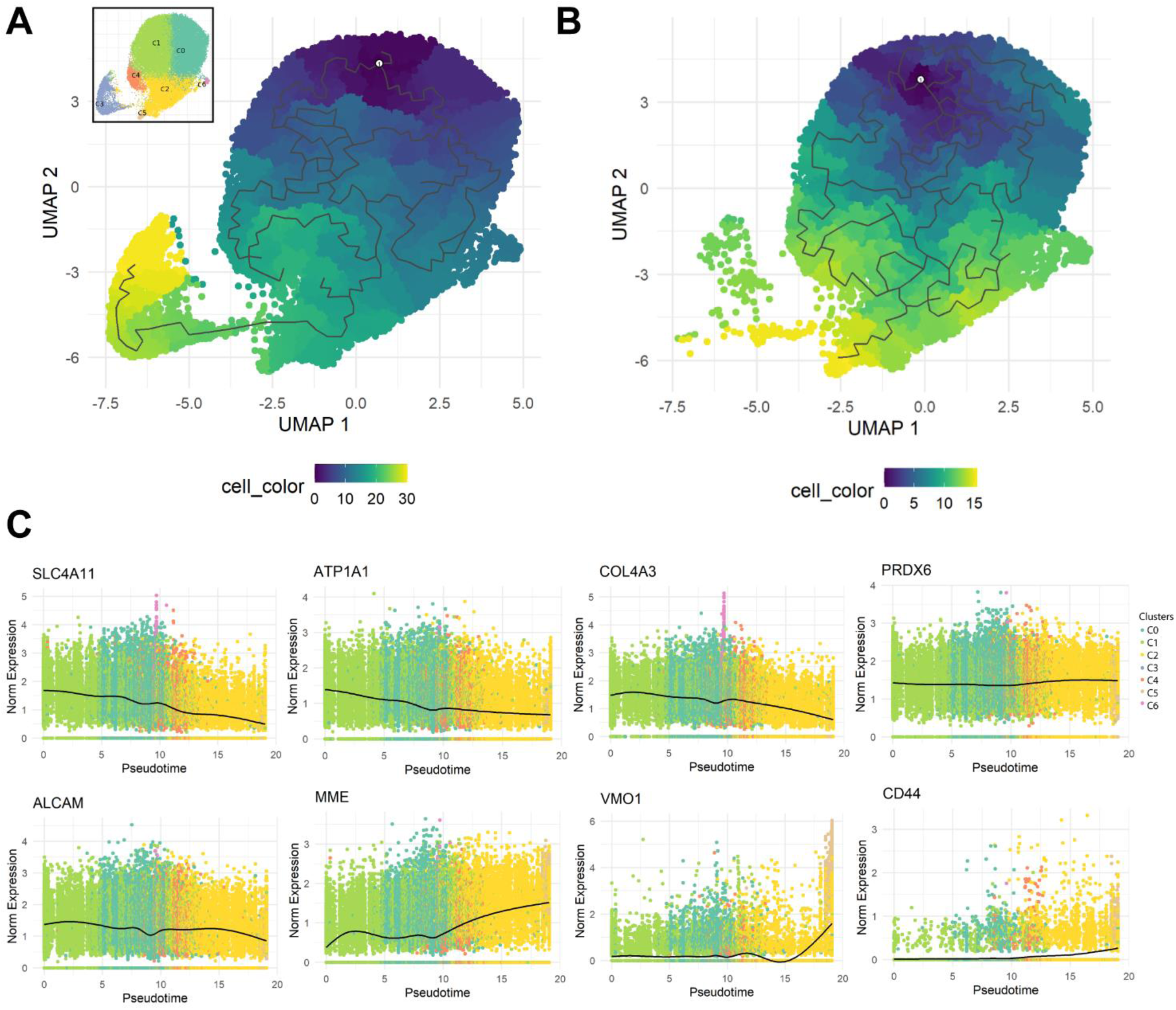
Pseudo temporal trajectory reconstruction reveals the dynamics of primary cultured CECs. (A) Monocle 3 pseudo temporal trajectory reconstruction on UMAP reduction of the scRNAseq confluency time points reveals the CEC cluster dynamics over primary culture. UMAP reduction is colored by pseudo time bins with dark blue being the earliest and yellow corresponding to late. (B) Monocle 3 pseudo temporal trajectory reconstruction on UMAP reduction of the scRNAseq confluency time points excluding proliferative cluster C3 reduces bias and reveals the temporal dynamics of the CEC clusters at confluence level. UMAP reduction is colored by pseudo time bins with dark blue being the earliest and yellow corresponding to late. (C) Pseudo time reconstruction reveals differential gene expression trends over CEC clusters. Our analysis revealed a reduction over time of CEC markers *SLC4A11*, *ATP1A1*, *COL4A3*, and *ALCAM*, while *PRDX6* expression was constant. Conversely, *CD44*, *MME*, and *VMO1* expression significantly increased over time.

The pseudo time reconstruction revealed that the expression of functional markers such as *SLC4A11* and *ATP1A1* was reduced over time, showing that senescent (cluster C2 and C4) and fibrotic (cluster C5) cells had a highly reduced expression of crucial functional markers (Figure 5C). Moreover, pseudo time reconstruction also revealed reduced expression of *COL4A3* and *ALCAM* over time (Figure 5C). Interestingly, the expression of *PRDX6*, a known marker of CECs, remained constant and did not decrease over time in the senescent (C2 and C4) and fibrotic (C5) clusters (Figure 5C). Besides crucial CEC markers, our analysis also revealed a significant increase of *CD44* expression over time in clusters C2 and C5 CECs. Furthermore, the expression of *MME* (CD10) and *VMO1* in the pseudo time analysis was increased in the senescent and fibrotic clusters C2 and C5 (Figure 5C). These genes were also differentially expressed in lower quality clusters C2 and C5, respectively, and can be candidates to assess quality of primary cultured CECs.

### scRNAseq transcriptomic profiles for quality assessment of primary cultured CEC correlate with protein level expression

The differential gene expression across CEC clusters and the pseudo time reconstruction both showed that the quality of the primary cultured CEC could be evaluated with a specific set of markers to differentiate therapy-grade CEC (clusters C0, C1, and C6), from lower quality CECs transitioning towards a senescent or fibrotic phenotype (clusters C2 and C5). The expression of *ALCAM* (CD166), *CGNL1* (cingulin-like protein 1), and *NCAM1* (CD56) were higher in the clusters comprising therapy-grade CECs (clusters C0 and C1), and lower in the cells in clusters C2 and C5 (Figure 6A). Additionally, the expression of *CD44*, *MME* (CD10), *VMO1*, and *THBS2* were higher in the fibrotic cells in cluster C5 and lower in the cells in clusters C0, C1, and C6 (Figure 6A), representing markers that could be used to exclude low quality CECs.

**Figure 6.**
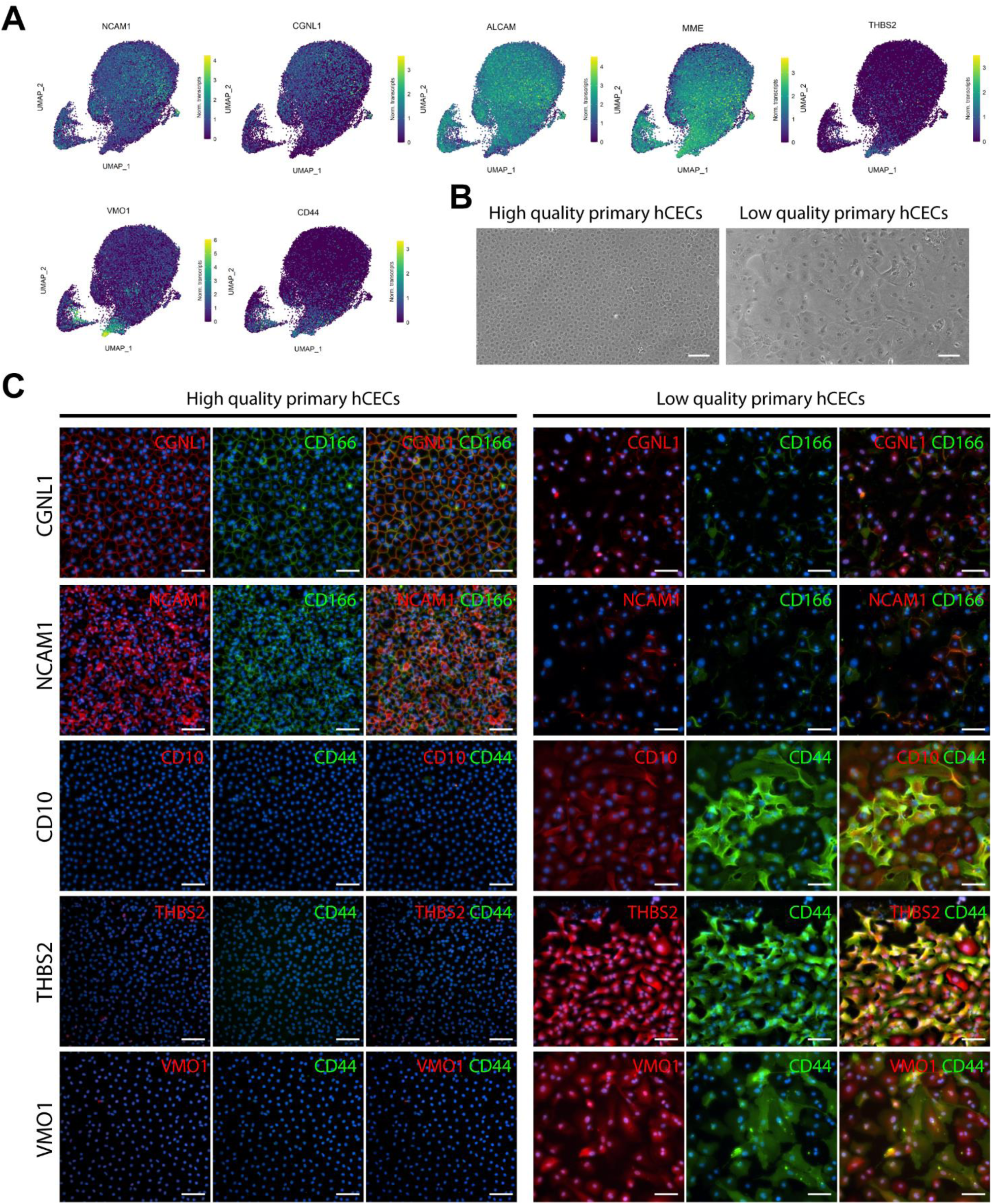
scRNAseq analysis suggests specific markers for CEC quality assessment. (A) Gene expression UMAP of differentially expressed markers in clusters of therapy-grade CECs (clusters C0 and C1) and clusters of senescent/fibrotic CECs (cluster C5). *NCAM1*, *CGNL1*, and *ALCAM* were differentially expressed in clusters CO and C1 (p < 0.01). *MME* and *CD44* were differentially expressed in clusters C2 and C5. *THBS2* and *VMO1* were differentially expressed in cluster C5 (p < 0.01). (B) Phase contrast images of a high quality therapy-grade CEC culture showing the typical hexagonal cell morphology and a low quality culture of primary human CECs showing the characteristic morphological alterations of an endothelial to mesenchymal transition. Scale bars represent 100 μm. (C) Immunofluorescence analysis shows expression of CD166 (green), NCAM1 (red), and CGNL (red) in high quality CEC cultures (N=2) and absence of expression in lower quality CEC cultures (N=2). Immunofluorescence analysis shows expression of CD44 (green), MME (CD10) (red), THBS2 (red), and VMO1 (red) in lower quality CEC cultures (N=2) and the absence of expression in high quality CEC cultures (N=2). Cell nuclei were stained with Hoechst 33342 (blue). Scale represent 100 μm.

To confirm the findings at the transcriptomic level were also present at the protein level, we assessed protein expression on primary CEC cultures with a characteristic hexagonal morphology associated with therapy-grade CECs and on primary CEC cultures composed of cells with spindle shape morphology, a characteristic of cells undergoing an endothelial to mesenchymal transition, referred as low quality primary CECs (Figure 6B). Immunofluorescence analysis confirmed that CGLN1, NCAM1 (CD56) and ALCAM (CD166) were exclusively expressed by good quality CECs and not expressed in the cultures containing CECs with an altered morphology (Figure 6C). Furthermore, immunofluorescence analysis also confirmed that CD44, MME (CD10), THBS2, and VMO1 were exclusively expressed by low quality CECs and not expressed in high quality CEC cultures (Figure 6C).

## DISCUSSION

In this study, we present a single-cell roadmap of human CECs in culture, revealing the diverse trajectories of individual cells. Our scRNAseq census of 42,200 primary cultured CECs revealed the presence of 7 clusters in therapeutically relevant time points, including therapy-grade CECs expressing *SLC4A11*, *ALCAM*, and *COL4A3* (clusters C0, C1 and C6); highly proliferative CECs expressing *MKI67* and *CENPF* (cluster C3); lower quality CECs entering senescence and EMT expressing *CDKN1A*, *CDKN2A*, and *TAGLN* (clusters C2 and C4); as well as fibrotic CECs expressing *CD44*, and *ACTA2* (cluster C5). We assessed to which extent CECs in culture resemble native human CECs and analyzed how these CEC populations diverge over culture time, giving insights into the alterations arising during primary culture. Moreover, our transcriptomic profiling provides an array of combinatorial markers to differentiate therapy-grade CEC from cells undergoing senescence and EMT, thereby paving the way for improving culture protocols and guiding the selection of cells for therapy. The transcriptomic data we obtained will help better our understanding of the mechanisms involved in the alterations occurring during primary culture of CECs leading to a loss of function and phenotype.

Our analysis showed that proliferating sub-confluent CECs sequenced at day 2 and day 5 clustered together (cluster 3), with high differential expression of ribosomal-related genes. We hypothesize that this is most likely due to their necessary adaptation to *in vitro* culture conditions and the use of proliferation media, which biased the first clustering. These findings led us to separately explore the 37,158 cells at confluency in passages 0, 1, and 2. Our analysis revealed three clusters of therapy-grade CEC (clusters C0, C1, and C6) based on the high differential expression of functional CEC markers *SLC4A11* and *ATP1A1* and the CEC markers *ALCAM*, *PRDX6*, and *COL4A3.* These cells were the majority in all passages, comprising 70% of all the sequenced cells. GSEA revealed that cells in C0 and C1 were metabolically active with increased expression of genes related to oxidative phosphorylation, in line with the crucial activity and function of CECs (Deguchi et al., 2022).

The integration analysis of the dataset from this study with native CECs from a previously published cornea cell atlas (Catala et al. 2021) revealed that native CECs clustered close to clusters C0, C1 and C6, suggesting the similarity of these cells. Cluster prediction analysis on native CECs revealed that these cells would group within clusters C1 (53.9%), C6 (45.1%), and C0 (0.4%). The low 0.4% prediction for cluster C0 is interesting because the primary cultured cells still expressed high levels of endothelial markers, namely *SLC4A11*, *ATP1A1*, *ALCAM*, and *PRDX6*. We hypothesize the low prediction might be due to a slight increase in ribosomal protein expression or a difference in the number of genes/cell which can be a technical sampling variation compared to clusters C1 and C6, which skews the cell clustering. Overall, our data shows that CECs in clusters C0, C1, and C6 are good quality CECs and are a suitable source for therapy. And while our dataset shows that cells in clusters C1 and C6 more closely resemble native human CECs that cells in cluster C0, it does not mean that cells in cluster C0 are unsuitable for therapy, but that they are distinguishable from native corneal endothelium.

Our scRNAseq analysis also revealed the presence of two clusters composed of senescent cells that were transitioning towards a mesenchymal phenotype (clusters C2 and C4). Our pseudo time reconstruction showed these cells were originating from the therapy-grade clusters C0, C1, and C6. This finding shows the transition to senescence of CECs during primary culture. Senescent cells in cluster C2 had reduced expression of key functional markers such as *SLC4A11*, *ATP1A1*, and *ALCAM*. This finding is in line with a recently published report that demonstrated a decrease of *SLC4A11* in lower-quality primary CECs (Deguchi et al., 2022).

The total number of senescent cells (clusters C2 and C4) increased over culture time, suggesting the CECs transitioned towards senescence over extended culture times. Cells comprising cluster C4 decreased over culture time, indicating that they either transition into senescent cells in cluster C2 or represent an end-point cluster, where cells tend to die over time. GSEA revealed that cells in cluster C2 had differential gene expression, specifically an increase in genes involved in the p53 and Rho GTPase pathways, suggesting these pathways might play a key role on the senescence and endothelial to mesenchymal transition of primary cultured CECs. P53 is a known senescence regulator (Mijit et al., 2020; Rufini et al., 2013), and its inhibition could delay the cellular senescence in primary CECs. A study in 2013 revealed that the inhibition of p53 was associated with improved morphology and higher expression of CEC markers, namely collagen type 8, Na/K ATPase, and N-cadherin, in primary cultured CECs (Sha et al., 2013). Furthermore, our findings suggest that the inhibition of the Rho GTPase pathway can play a key role in delaying cellular senescence, further confirming that the use of Rho-associated protein kinase (ROCK) inhibitors such as Y-27632 might be a pivotal factor in the protocols for primary expansion of therapy-grade CECs. Indeed, previous studies have used Y-27632 for the primary expansion of CECs (Kinoshita et al., 2018; Okumura et al., 2009; Parekh et al., 2020; Peh et al., 2015b). Based on these findings, we therefore recommend the use of ROCK inhibitors during the primary expansion of CECs.

Our scRNAseq analysis revealed that a cluster of CECs (cluster C5) expressing characteristic fibrotic markers ACTA2 and CD44 originated from the senescent cells at clusters C2, suggesting a transition from senescence to endothelial to mesenchymal transition phenotype. While fibrotic markers were found in later time points, we did observe that cells comprising cluster C5 appeared as early as passage 1, but were then reduced by passage 2. This finding could be due to the sequencing sampling limitation of 10,000 cells per sample, making it highly possible that this small fibrotic cell population was not sequenced from a culture of hundreds of thousands of cells. Our second hypothesis is that after passaging, the fibrotic cells could not successfully adhere, causing a reduction of this cell population and enriching for good quality cells.

Our findings showed that extended culture times decreased the proliferation potential of CECs, shown by a reduction in the number of cells in the proliferation cluster C3 across passages. Moreover, we detected the presence of a subpopulation of undesired proliferative fibrotic cells that could potentially overgrow the culture of CECs over extended culture periods. In our view, these results show that with the current protocols, culturing CECs further than passage 2 is incompatible with their therapeutic use, a recommendation in line with previous studies that suggested passage 2 as the threshold time point to assure the therapeutic suitability of primary cultured CECs (Frausto et al., 2016, 2020; Peh et al., 2019).

Selecting and assessing the quality of the primary cultured CECs are a crucial aspect to ensure a safe and efficacous therapy. Based on our differential expression analysis and pseudo time reconstruction, we show that therapy-grade CEC should be identified by the expression of CD166 and NCAM1 membrane proteins together with CGNL1, a membrane-associated protein to cellular tight junctions; lack of expression of altered extracellular matrix, namely VMO1 and THBS2; and lack of expression of membrane proteins CD44 and CD10. While the membrane protein markers suggested by our analysis would allow sorting for therapy-grade CECs, we believe is equally important to characterize CEC culture quality based on expression of other fundamental proteins such as aberrant extracellular matrix production. Future studies are required to understand how the expression of these markers correlate to therapeutic success. Similar to our suggestion to analyze markers for therapy-grade CECs (CD166^+^, NCAM1^+^, CGNL1^+^, CD44^-^, CD10^-^, VMO1^-^, and THBS2^-^), Kinoshita and colleagues proposed the combinatorial marker expression referred as the E-ratio (CD166^+^, CD44^-^, CD133^-^, CD24^-^, and CD105^-^) to assess for therapy-grade CECs. We and they both detected increased CD166 expression in therapy-grade CECs and CD44 exclusively expressed in lower quality senescent and transitioning CECs. By contrast, our analysis revealed that *CD24* and *ENG* (CD105) were heterogeneously and minimally expressed across clusters in some CECs, and we did not detect expression of *PROM1* (CD133) in any cluster. These differences might be due to the lack of correlation between transcript and protein detection. Future studies analyzing such differences can shed light on the suitability of markers to assess or enrich for therapy-grade CECs.

While primary cultured CECs have been traditionally assessed as bulk entities without accounting for their heterogeneity, our study analyses them at the single-cell level over five culture time points in three different passages. Our study provides significant information to help understand the changes arising from the culture of human CECs, portraying their cellular heterogeneity as well as characterizing their variability over extended culture times. These results provide a pivotal dataset that can help identify and characterize the undesired cell populations arising from primary culture in the attempt to improve current protocols. Our results also show the importance of supplementing media for CEC expansion with ROCK inhibitors to reduce cellular senescence. Furthermore, based on the results reported in this study, we propose a combination of markers to assess the quality of primary cultured CECs. Overall, this transcriptomic cell analysis offers a baseline for future studies with the aim of improving CEC- based therapies.

## ACKNOWLEDGEMENTS

The authors thank Single Cell Discoveries (Utrecht, the Netherlands) for the single cell sequencing services provided. The authors thank the Lions Eye Institute for Transplant & Research (Tampa, FL, USA) for providing research-grade human corneas.

## AUTHOR CONTRIBUTIONS

Conceptualization: P.C., N.G., V.L.S.L., M.M.D.; Investigation: P.C., N.G.; Resources: P.C., N.G., V.L.S.L., M.M.D.; Data Curation: N.G.; Writing – Original Draft: P.C.; Writing – Review & Editing: N.G., V.L.S.L., M.M.D.; Supervision: V.L.S.L., M.M.D.; Funding acquisition: P.C., V.L.S.L., M.M.D. All authors ensured that questions on the accuracy or integrity of all parts of the study were appropriately researched and resolved.

## MATERIALS AND METHODS

### RESOURCE AVAILABILITY

#### Lead contact

Further information and requests for resources should be addressed to the lead contacts, Mor

M. Dickman or Vanessa L.S. LaPointe.

#### Materials availability

This study did not generate new reagents or materials

#### Data and code availability

The data regarding this study will be deposited to the Gene Expression Omnibus upon manuscript acceptance. The R code used for this study will be publically available at DataVerse.NL (including DOI) upon manuscript acceptance.

Additional information required to reanalyze the data reported in this paper is available from the lead contact upon request.

### EXPERIMENTAL MODEL AND SUBJECT DETAILS

#### Research-grade donor human corneas and ethical statement

This study was performed in compliance with the tenets of the Declaration of Helsinki. All research-grade human donor corneas used for primary culture were obtained from the Lions Eye Institute for Transplant & Research (Tampa, USA), with informed consent from the next of kin. The research involving human-derived corneas was performed in accordance to Maastricht University and Dutch national regulations. All corneas had an endothelial cell density of at least 2800 cells/mm^2^, deemed unsuitable for transplantation, and were preserved in Optisol-GS at 4°C for up to 14 days prior to their use (Table 1).

### METHOD DETAILS

#### Isolation and culture of primary human corneal endothelial cells

Six paired corneas, from two male and one female donor aged 27 to 34 years, were used for isolation and primary culture of endothelial cells. Donors had no history of ocular disease, chronic systemic disease, or pathological infection such as HIV, Hepatitis B and C, HTLV-I/II, syphilis, or SARS-CoV-2.

Prior to isolation, the endothelial–Descemet’s corneal layer was manually stripped as follows: the corneas were vacuum fixed in a punch base (e.janach) endothelial cell side up and trephined with a 10 mm Ø corneal punch at a fixed depth of 100 μm (e.janach). To delimit the endothelial trephined line, corneas were stained with a trypan blue solution (0.4%) for 30 s, and washed with balanced salt solution sterile irrigating solution (BSS; Alcon). The corneal endothelium was then gently lifted using a DMEK cleavage hook (e.janach) and fully stripped using angled McPherson tying forceps.

Human corneal endothelial cells were isolated and cultured as previously reported (Peh et al., 2015a). Briefly, the stripped endothelium–Descemet’s layer was incubated with 2 mg/ml collagenase A (Roche) solution in human endothelial serum free media (SFM) (Thermo Fisher Scientific) for 2–5 h at 37°C followed by a 5 min incubation in TrypLE express (Thermo Fisher Scientific) to generate small clumps of corneal endothelial cells. Cells from each cornea were seeded equally across 2 wells of a 24-well plate coated with fibronectin collagen (FNC) coating mix (Athena Enzyme Systems) in M5 stabilization media (human endothelial SFM (Thermo Fisher Scientific) supplemented with 5% fetal bovine serum (FBS), 100 U/mL penicillin– streptomycin, and 0.25 μg/mL amphotericin B) supplemented with 10 μM Y-27632 (STEMCELL Technologies). Subsequently, corneal endothelial cells were cultured in M4 proliferation medium (1:1 Ham’s F12 (Thermo Fisher Scientific) and M199 (Thermo Fisher Scientific) supplemented with 5% FBS, 20 μg/mL ascorbic acid (Sigma), 1 × ITS (Thermo Fisher Scientific), 10 ng/mL human recombinant bFGF (Sigma), and 10 μM Y-27632 (STEMCELL Technologies)); media was refreshed every other day. Upon reaching 90% confluency, after approximately 8–10 days of culture, cells were cultured in M5 stabilization media for 7 days. After this, corneal endothelial cells were treated with TripLE express and passaged into wells pre-coated with FNC-coating mix at a seeding density of 10,500 cells/cm^2^ in M5 stabilization medium. All cell culture was performed in incubators with humidified atmosphere of 37°C and 5% CO2.

### Preparation of single cell suspension and methanol fixation

Cells from five different culture time points were methanol fixed for sequencing. Namely, cells at days 2 and 5 of culture after isolation in M4 proliferation media at passage 0, and cells at confluency after 7 days of culture in M5 stabilization media, at passages 0, 1, and 2.

To generate a single cell suspension, primary cultured corneal endothelial cells were treated with TripLE express for approximately 30 min at 37°C. Then cells were centrifuged for 5 min at 800 × *g* and resuspended in 1 mL ice-cold Dulbecco’s phosphate-buffered saline (DPBS). Next, the cells were centrifuged for 5 min at 800 × *g* and resuspended in ice-cold DPBS at a ratio of 200 μL DPBS/1 × 10^6^ cells, followed by the dropwise addition of ice-cold methanol at a ratio of 800 μL DPBS/ 1 × 10^6^ cells. The fixed cell suspensions were stored at -80°C until sequencing.

### Single-cell RNA sequencing

scRNAseq of primary cultured CECs was performed at Single Cell Discoveries (Utrecht, the Netherlands) following standard 10× Genomics 3’ V3.1 chemistry protocol. Cells were rehydrated and loaded on the 10× Chromium controller as follows. Approximately 10,000 cells were loaded per each sample specified in Table 2. The resulting sequencing libraries were prepared following a standard 10× Genomics protocol and sequenced with an Illumina NovaSeq 6000 platform; read length: 150 bp, paired-end.

### Bioinformatic analysis of scRNA-seq data

The BCL files resulting from sequencing were transformed to FASTQ files with 10× Genomics Cell Ranger mkfastq following its mapping with Cell Ranger count. During sequencing, Read 1 was assigned 28 bp, and were used for identification of the Illumina library barcode, cell barcode and unique molecular identifier (UMI). R2 was used to map the human reference genome GRCh38. Filtering of empty barcodes was done in Cell Ranger. The data from all samples were loaded in R (version 4.2.0) (R Core Team, 2013) and processed using the Seurat package (version 4.1.1) (Stuart et al., 2019). More specifically, for each library a UMI cutoff was used to filter out low quality cells because of the differences between the libraries (*i.e.* g1 – 500, g2 – 3000, g3 – 500, g4 – 1313, g5 – 4000, g6 – 4000, g7 – 4000, g8 – 4000, g9 – 4000) (Table 2). Additionally, cells with less than 10% mitochondrial gene content were retained for analysis. The data of all 10× libraries were merged and processed together. The merged dataset was normalized for sequencing depth per cell and log-transformed using a scaling factor of 10,000. The multiplexed samples were demultiplexed based on their snp profile using Souporcell (Heaton et al., 2020). Briefly, the bam file and barcodes of each library were used as input together with the reference genome GRCh38. Besides the default parameters, the number of clusters was set to the number of multiplexed samples per library. The demultiplexing information for each cell was added to the metadata object in Seurat. The patient and library effect was corrected using Harmony (Korsunsky et al., 2019), as implemented in Seurat and used for dimensionality reduction and clustering of all cells. Cells were clustered using graph-based clustering and the original Louvain algorithm was utilized for modularity optimization. The differentially expressed genes per cluster were calculated using the Wilcoxon rank sum test and used to identify cell types. Putative doublets were computationally identified using scDblFinder (version 1.2.0) (Germain, 2020) but did not compose a separate cluster and therefore were not removed from the dataset (Figure S9). Pseudo time analysis was performed using the Monocle-3 package (version 1.0.0) (Cao et al., 2019). Gene set enrichment analysis was performed on lists of differentially regulated genes without prefiltering step. Gene lists were preranked using the signed -log10 (P adj) and subjected to enrichment analysis using fgsea package (version 1.22.0) (Korotkevich et al., 2021) and gage (version 2.46.1) (Luo et al., 2009) with curated and hallmark gene sets from MSigDB Collections (version 7.5.1) (Liberzon et al., 2015; Subramanian et al., 2005). To prune selectively the resulting pathways and GO terms, enrichment was considered when up- or downregulated gene sets were detected using both methods. The CEC dataset was integrated with the previously published cornea atlas (Català et al., 2021). The library effect was corrected for using harmony, followed by dimensionality reduction. The cluster information from the separate analysis was used to overlay in 2D space. The cell type of the atlas cells was predicted using the CEC confluency dataset using the cell label transfer functionality from Seurat.

### Immunofluorescence

Primary cultured CECs at time point confluency passage 2 deriving from all three donors were used for immunofluorescence analysis. CECs were fixed in 4% PFA for 15 min at ambient temperature and the cells were permeabilized with 0.1% (v/v) Triton X-100 in phosphate buffered saline (PBS) for 10 min. After permeabilization, non-specific antibody interactions were blocked with blocking buffer (2% (w/v) BSA solution in PBS) for 1 h at ambient temperature. CECs were incubated overnight at 4**°**C with primary antibodies mouse monoclonal anti-CD166 [3A6] (1:200 dilution, BD Biosciences), rabbit polyclonal anti-ZO1 (1:100 dilution, Thermo Fisher Scientific), mouse monoclonal anti-CD44 [Hermes-3] (1:400 dilution, Abcam), rabbit polyclonal anti-VMO1 (1:100 dilution, Prestige Antibodies), rabbit polyclonal anti-THBS2 (1:100 dilution, Abcam), rabbit monoclonal anti-CD10 [EPR22867-118] (1:100 dilution, Abcam), rabbit monoclonal anti-NCAM1 [CAL53] (1:100 dilution, Abcam), and rabbit polyclonal anti-CGNL1 (1:200 dilution, Atlas antibodies) diluted in blocking buffer. After primary antibody incubation, tissues were washed three times in PBS and then incubated with secondary antibodies goat anti-mouse A488 (1:400 dilution; Thermo Fisher Scientific), and donkey anti-rabbit A568 (1:400 dilution; Thermo Fisher Scientific) diluted in blocking buffer for 50 min at ambient temperature in the dark. Cell nuclei were stained with 1 μg/mL Hoechst 33342 (Thermo Fisher Scientific) for 10 min. The CEC samples were then washed three times in PBS and examined on an Eclipse Ti-E inverted microscope (Nikon) equipped with an X-Light V2-TP spinning disk (Crest Optics).

### STATISTICAL ANALYSIS

#### Statistical and quantitative analysis of scRNAseq data

All the statistical analysis for scRNAseq were performed in R (version 4.2.0) with the packages described in the methods detail section. Briefly, differentially expressed genes were detected using a Wilcoxon Rank-Sum test, statistical significance was defined as p < 0.01. GSEA revealed differentially enriched gene sets from MSigDB Collections, statistical significance was defined as p < 0.05.

## SUPPLEMENTAL INFORMATION TITLES AND LEGENDS

**Figure S1.**
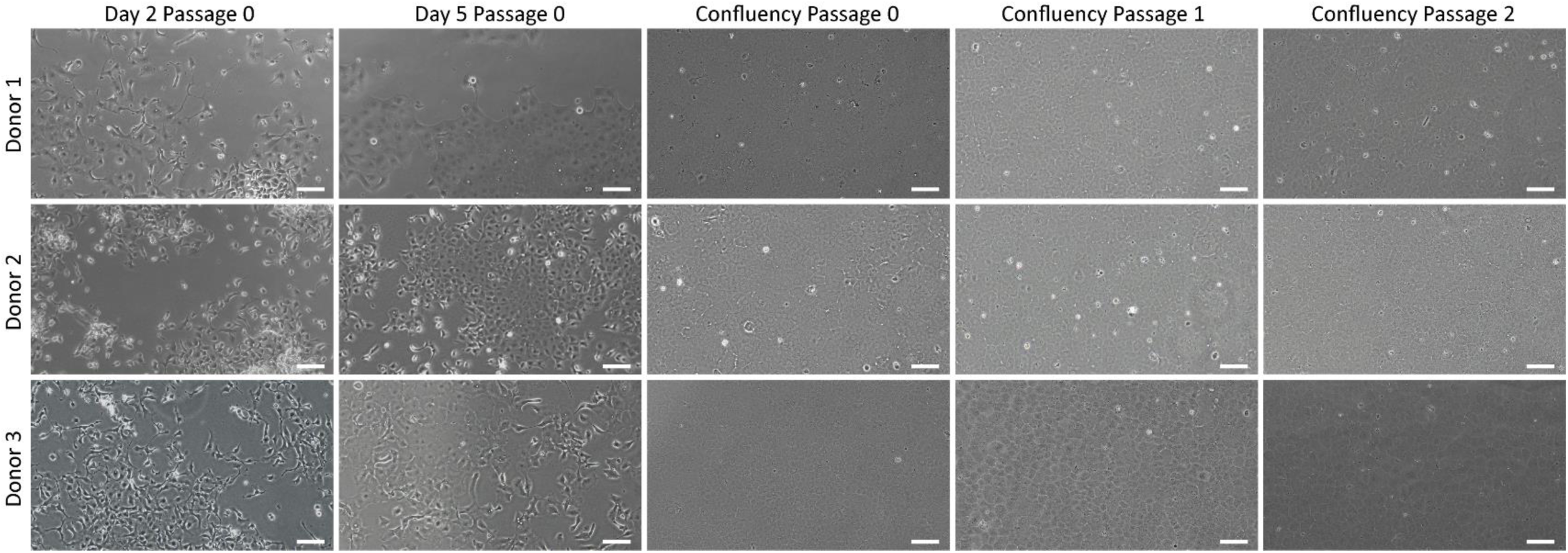
Phase contrast images of each donor time point used for scRNAseq, related to Figure 1. Phase contrast imaging shows desired endothelial morphology in all of the sequenced samples. Scale bar represents 100 μm.

**Figure S2.**
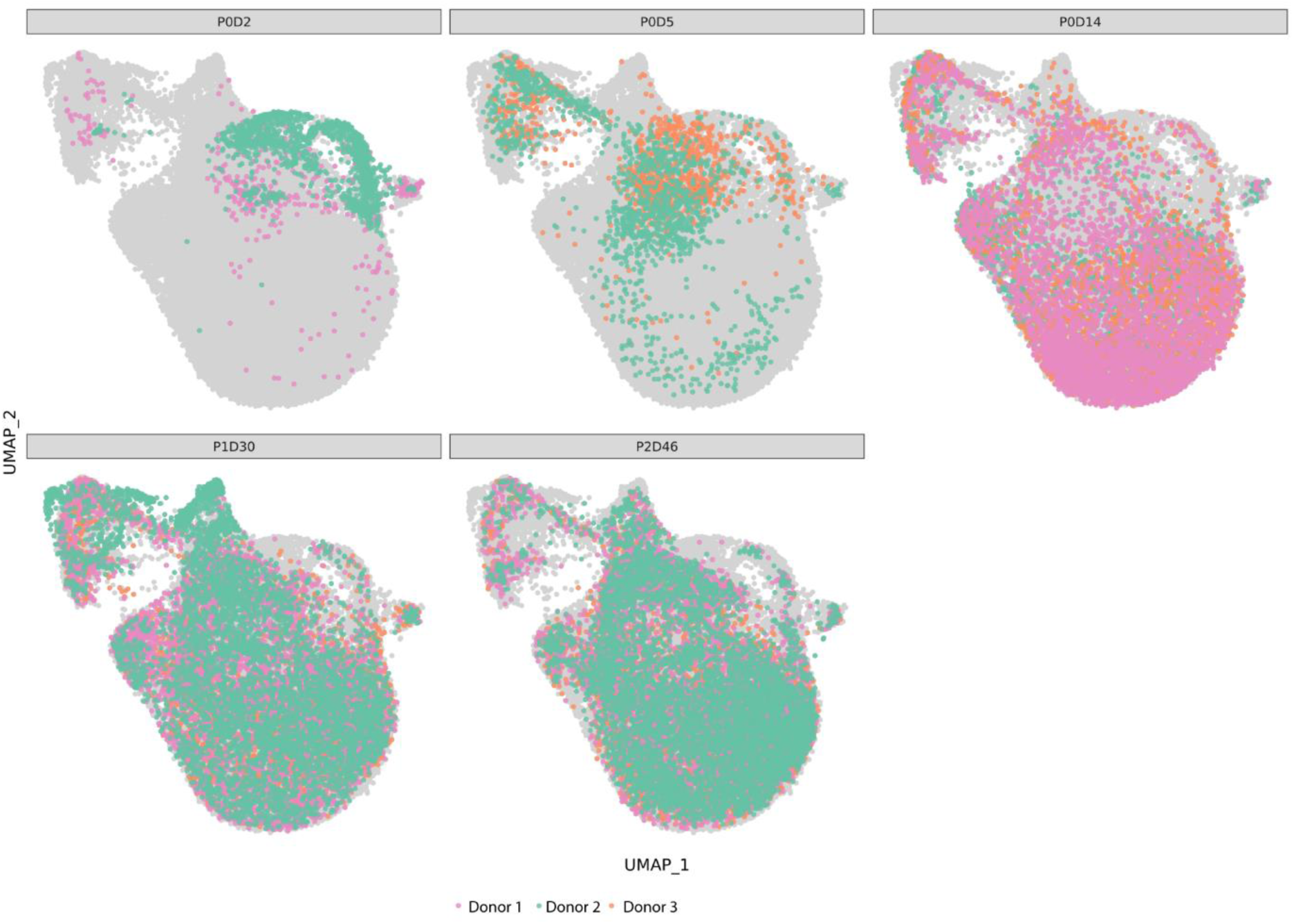
UMAP projection of cells per time point, related to Figure 2. Cell distribution UMAP per each time point further confirms that cluster is composed of CEC at early culture time points. Each donor is represented by color confirming the donor distribution across cell clusters. P0D2: day 2 in proliferation media. P0D5: day 5 in proliferation media. P0D14: passage 0 maintenance media. P1D30: passage 1 maintenance media. P2D46: passage 2 maintenance media.

**Figure S3.**
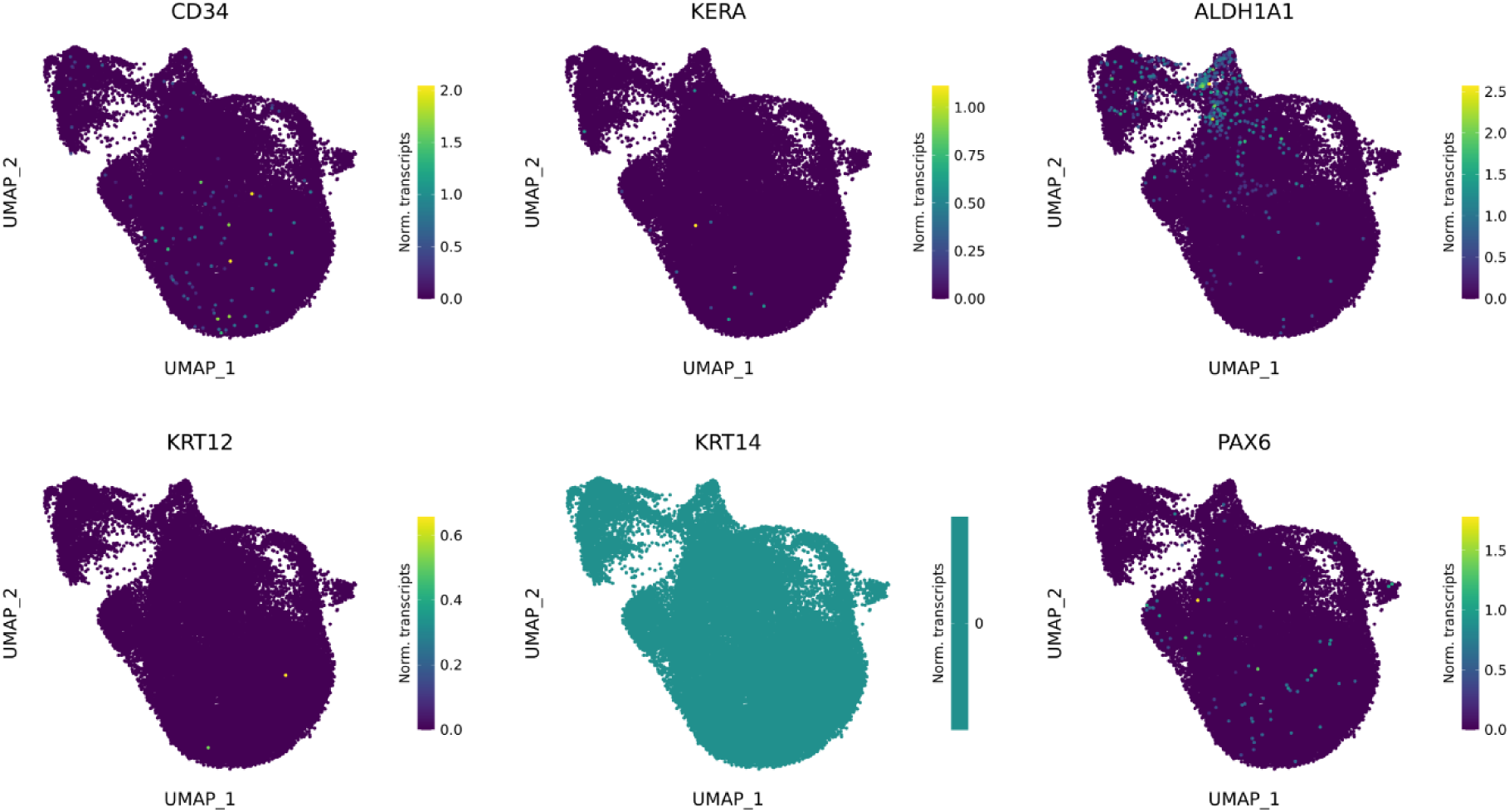
scRNAseq revealed absence of stroma and epithelial contamination, related to Figure 2. Gene expression UMAP of stromal markers *KERA*, *LUM*, and *ALDH1A1* and epithelial markers *KRT12*, *KRT14*, and *PAX6* confirmed absence of contaminant corneal side- populations in the primary culture.

**Figure S4.**
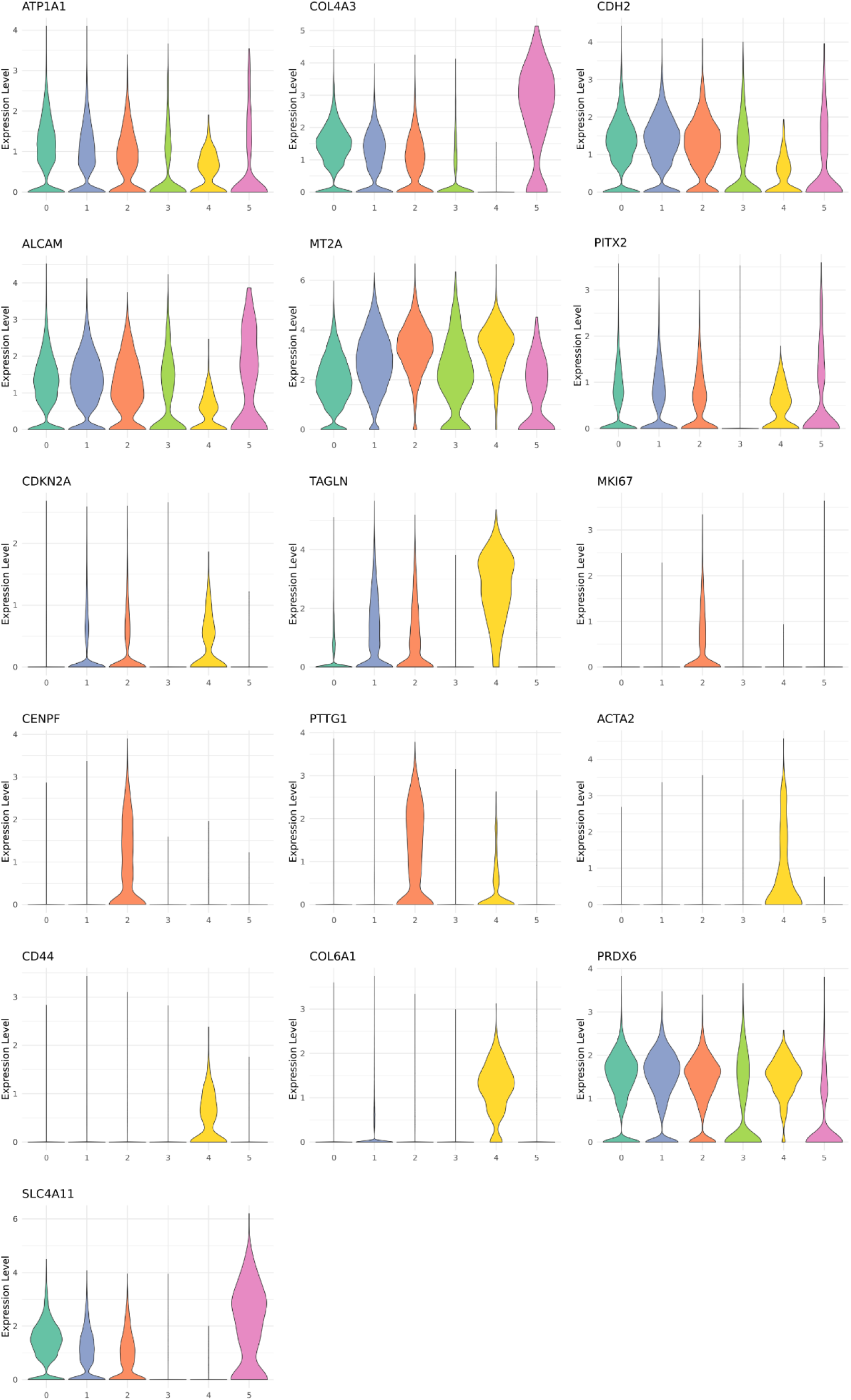
Violin plots of differentially expressed genes for cluster identification, related to Figure 2. Violin plots of differentially expressed genes for cluster annotation. Corneal endothelium: *AP1A1, COL4A3, CHD2, ALCAM, SLC4A11, PITX2*. Senescence: *MT2A, CDKN2A, TAGLN*. Proliferation: *MKI67, CENPF, PTTG1*. Fibrosis and endothelial to mesenchymal transition: *ACTA2, CD44, COL6A1*.

**Figure S5.**
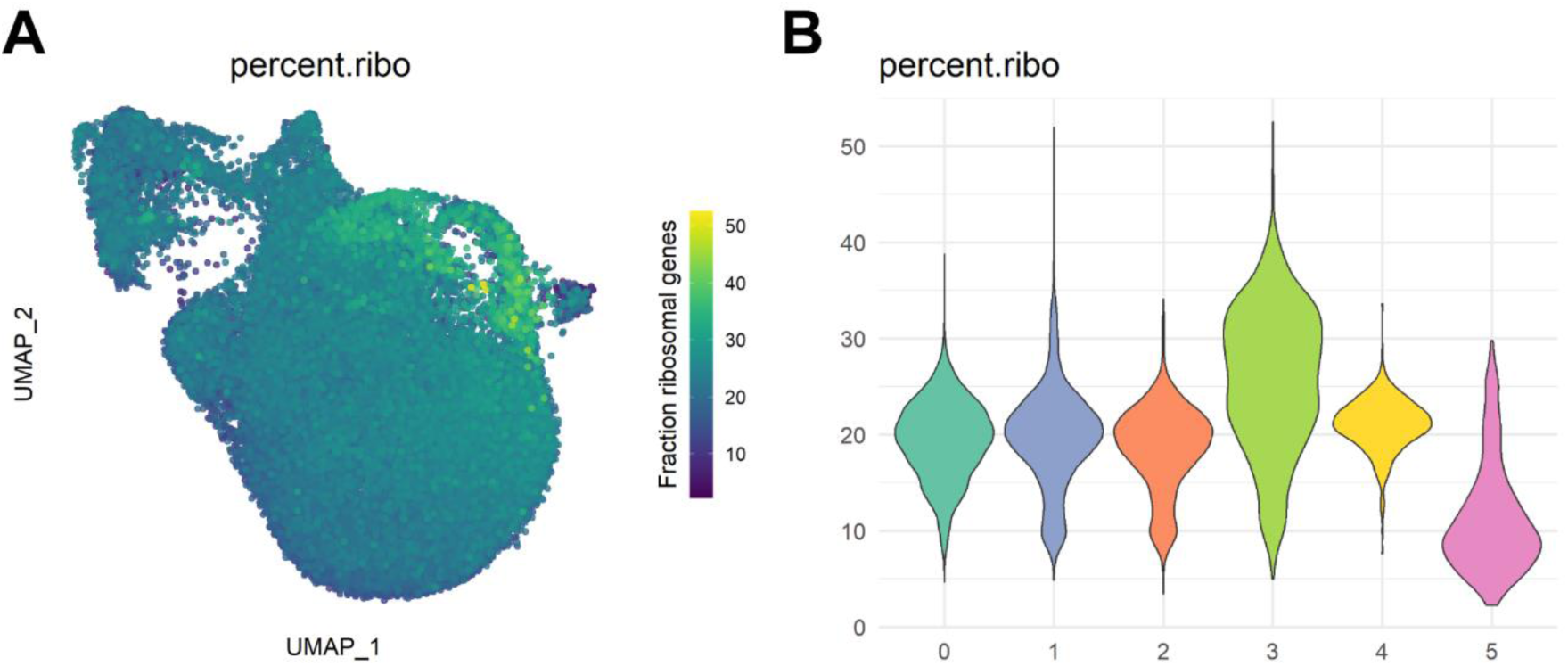
Cluster 3 is enriched in ribosomal gene expression. (A) UMAP representation of ribosomal gene fraction shows that cluster 3 expresses a high amount of ribosomal genes compared to other clusters. (B) Violin plot of ribosomal gene fraction shows further confirms a high expression of ribosomal genes in cluster 3 compared to the other detected clusters.

**Figure S6.**
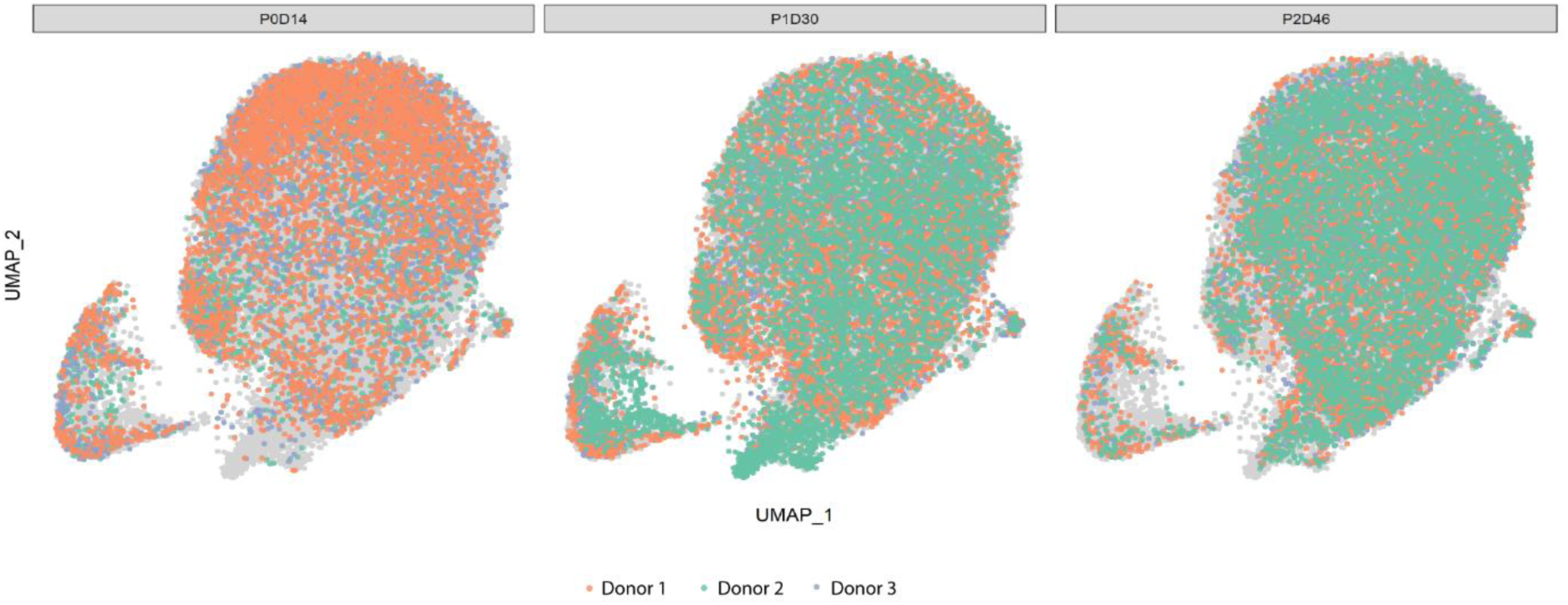
UMAP projection of cells per time point at confluency, related to Figure 3. Cell distribution UMAP per each time point at confluency level shows homogeneous distribution of donors across all cell clusters. P0D14: passage 0 maintenance media. P1D30: passage 1 maintenance media. P2D46: passage 2 maintenance media.

**Figure S7.**
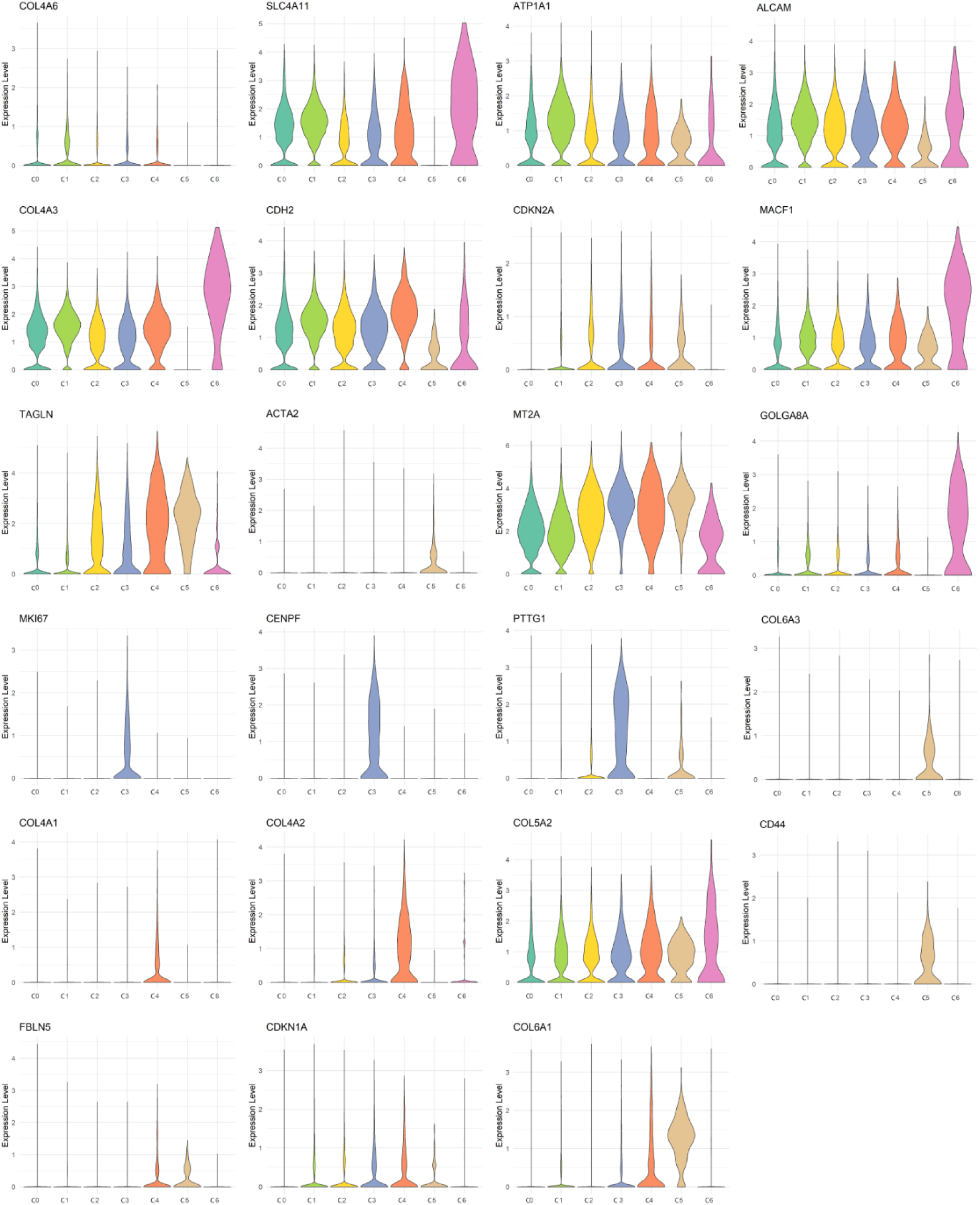
Violin plots of differentially expressed genes for cluster identification, related to Figure 3. Violin plots of differentially expressed genes for cluster annotation at the confluency time points. Corneal endothelium: *SLC4A11*, *ATP1A1*, *ALCAM*, *CDH2*. Corneal endothelium extracellular matrix: *COL4A8*, *COL4A3*, *COL4A1*, *COL4A2*, *COL5A2*. Senescence: *CDKN2A*, *TAGLN*, *MT2A*, *CDKN1A*, *LGALS1*. Cell secretion: *GOLGA8A*. Proliferation: *MKI67*, *CENPF*, *PTTG1*. Fibrosis and endothelial to mesenchymal transition: *COL6A3*, *CD44*, *FBLN5*, *COL6A1*, *ACTA2*

**Figure S8.**
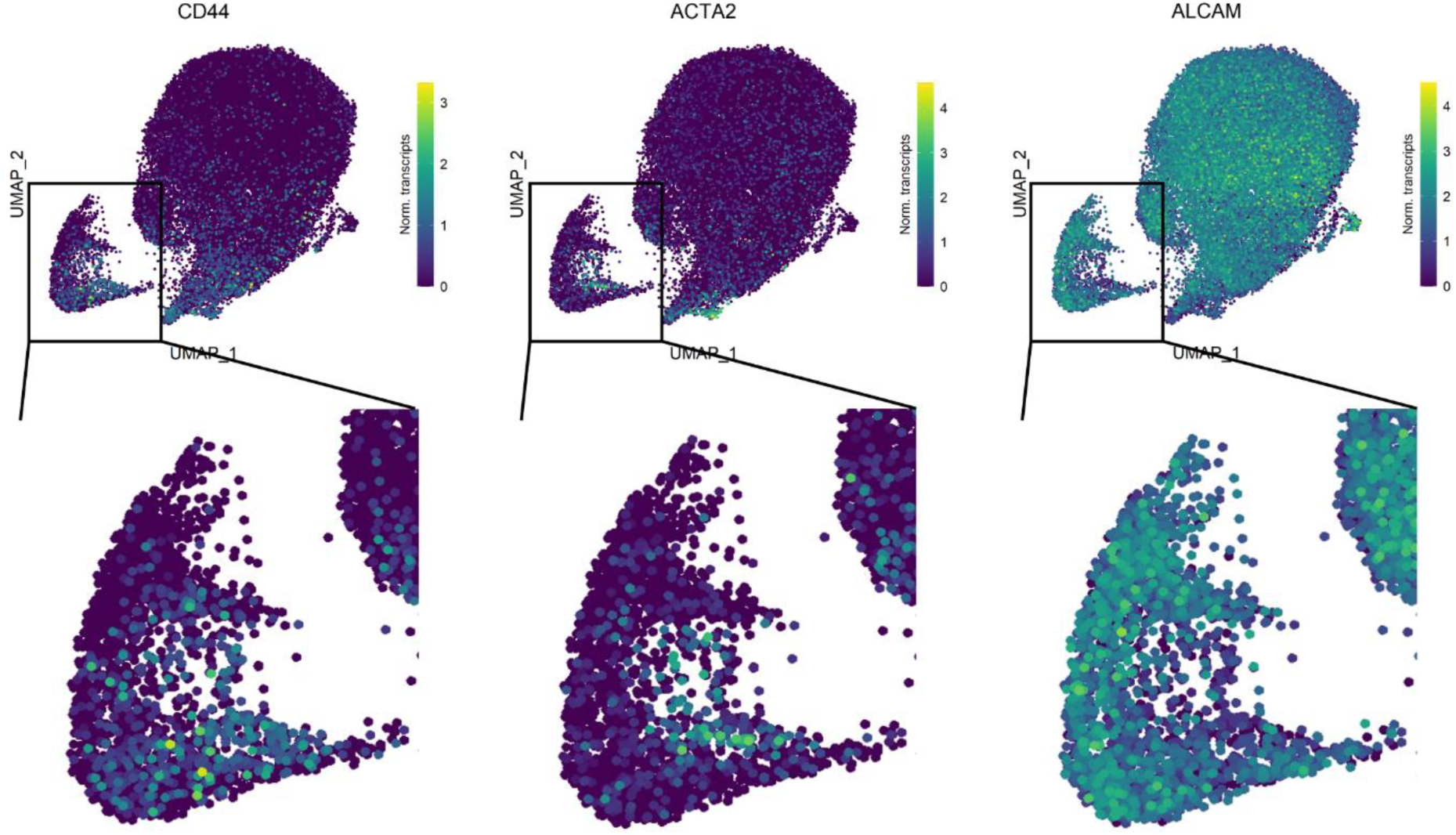
Gene expression UMAP of *ALCAM* (CD166), *CD44* and *ACTA2*, related to Figure 3. Gene expression UMAP shows heterogeneous expression of *ALCAM*, *CD44* and *ACTA2* across cell cluster C3.

**Figure S9.**
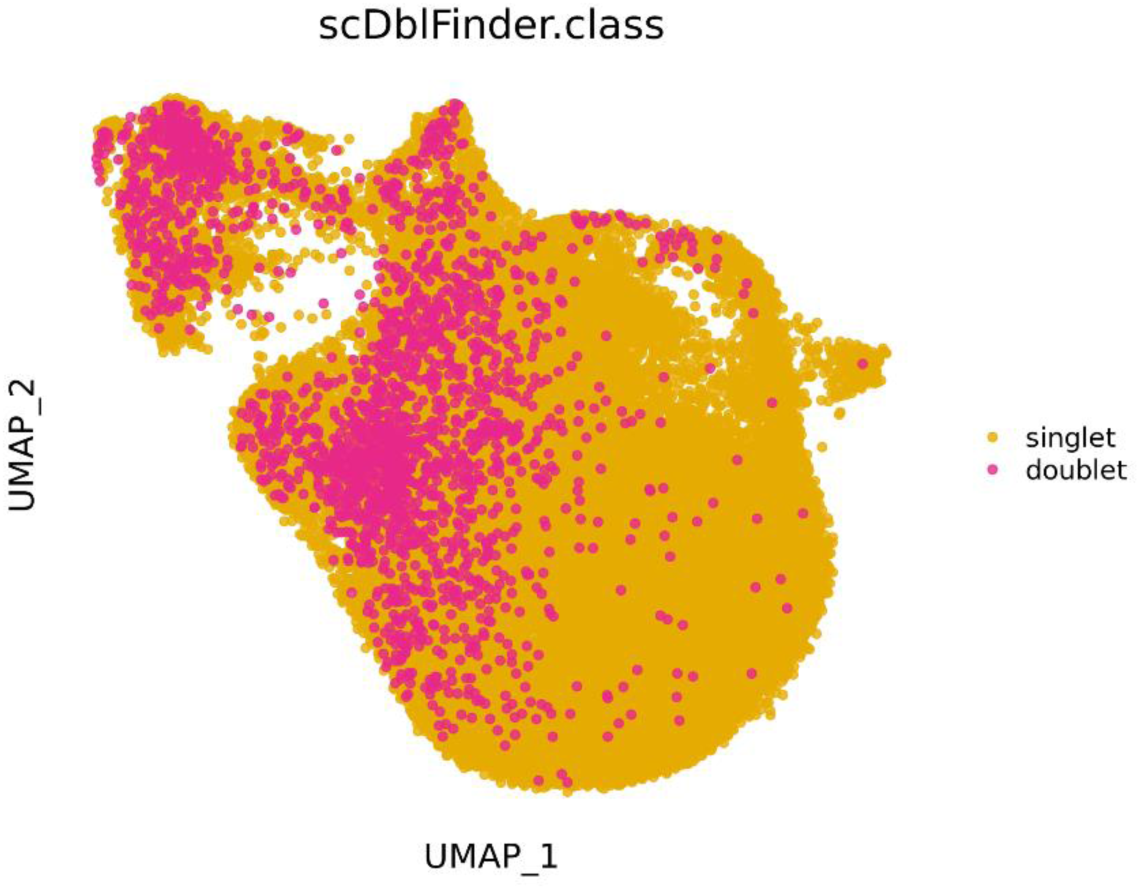
Identification of putative doublets. Putative doublets were identified with scDblFinder. The identified putative doublets were dispersed across all cell clusters and did not bias scRNAseq clustering therefore were not removed.

